# HiPLA: High-throughput imaging Proximity Ligation Assay

**DOI:** 10.1101/371062

**Authors:** Leonid A. Serebryannyy, Tom Misteli

## Abstract

Protein-protein interactions are essential for cellular structure and function. To delineate how the intricate assembly of protein interactions contribute to cellular processes in health and disease, new methodologies that are both highly sensitive and can be applied at large scale are needed. Here, we develop HiPLA (high-throughput imaging proximity ligation assay), a method that employs the antibody-based proximity ligation assay in a high-throughput imaging screening format to systematically probe protein interactomes. Using HiPLA, we probe the interaction of 60 proteins and associated PTMs with the nuclear lamina in a model of the premature aging disorder Hutchinson-Gilford progeria syndrome (HGPS). We identify a subset of proteins that differentially interact with the nuclear lamina in HGPS. In combination with quantitative indirect immunofluorescence, we find that the majority of differential interactions were accompanied by corresponding changes in expression of the interacting protein. Taken together, HiPLA offers a novel approach to probe cellular protein-protein interaction at a large scale and reveals mechanistic insights into the assembly of protein complexes.

## 1. Introduction

Elucidation of how proteins interact is central to understanding cellular function and structure. As protein identification technology has advanced, so has the realization that a protein’s interactome is highly complex and tightly regulated. Traditionally, biochemical approaches such as immunoprecipitation followed by mass spectroscopy or yeast-two hybrid screens have facilitated identification of protein interactors [1, 2]. However, these methods are commonly based on interaction frequencies that occur outside of their native environment in the intact cell and they do not always translate to be biologically relevant [3]. Recently, labeling methods such as biotin ligase-based BioID [4, 5], ascorbate peroxidase-based APEX [6], or reversible chemical crosslinkers [7] have been employed to label interacting proteins in the cell, effectively capturing snapshots of native protein complexes. To complement these approaches, increased utilization of super resolution imaging, Förster resonance energy transfer (FRET), fluorescence correlation spectroscopy (FCS), protein fragment complementation assays (PCA), and proximity ligation assays (PLA) has enabled systematic probing and quantification of protein complex formation with nanometer spatial resolution *in vivo* [8]. Nevertheless, while fluorescent imaging is an ideal technique to probe protein-protein interactions in their natural context, these methods are largely low-throughput and do not allow for the large-scale interrogation of interactomes [8, 9].

A notable example for the importance of systematically delineating protein-protein interactions is Hutchinson-Gilford progeria syndrome (HGPS) [10-13]. HGPS is a rare premature aging disorder caused by activation of an alternative splice site in the *LMNA* gene which encodes the two nuclear intermediate filament proteins lamin A and C. Aberrant splicing of *LMNA* mRNA results in a 50 amino acid C-terminal deletion in the lamin A protein and the mutant proteins is referred to as progerin [14-18]. Unlike wild type lamin A, progerin remains farnesylated and tethered to the inner nuclear membrane. Progerin expression leads to the dysfunction of multiple tissues throughout the body, most prominently cardiovascular failure which is ultimately fatal due to myocardial infarction or stroke [19]. Proteomic comparisons demonstrate that progerin and lamin A have distinct interactomes [20-22], suggesting differential protein complex formation is involved in pathogenesis. Indeed, these interactions and others have been independently verified to be affected in HGPS using combined biochemical and imaging approaches [10-13]. However, the progerin interactome, particularly with lamin-associated partners, remains poorly characterized.

Given the current limitations in identifying and validating protein interactomes, we sought a means to screen the effects of cellular perturbations on specific protein-protein interactions in a high-throughput manner. Based on the ability of PLA to detect protein-protein interactions by means of imaging, we developed a high-throughput version of PLA. The PLA method utilizes primary antibodies to identify proteins of interest, similar to standard immunofluorescence (IF) labeling, however the secondary antibodies are tagged with short DNA strands instead of fluorophores [23] (**Figure 1**). If the two DNA strands are within ~40 nm, they are ligated and rolling circle amplification with fluorescently-conjugated nucleotides can be used to visualize interaction sites. PLA has been successfully adapted for a range of applications including to probe the interaction of proteins across cellular compartments, to map post-translational modifications (PTMs), to quantify protein amounts in solution, and to screen compounds for their kinase inhibitory activity [24, 25]. Here, we describe the methodology for high-throughput imaging proximity ligation assays (HiPLA) as a means to simultaneously test the effects of a cellular intervention on a large number of protein interactions. As proof-of-concept, we assay a library of antibodies against nuclear proteins in an inducible cellular model of HGPS to examine how progerin expression affects the lamin interactome.

**Figure 1:**
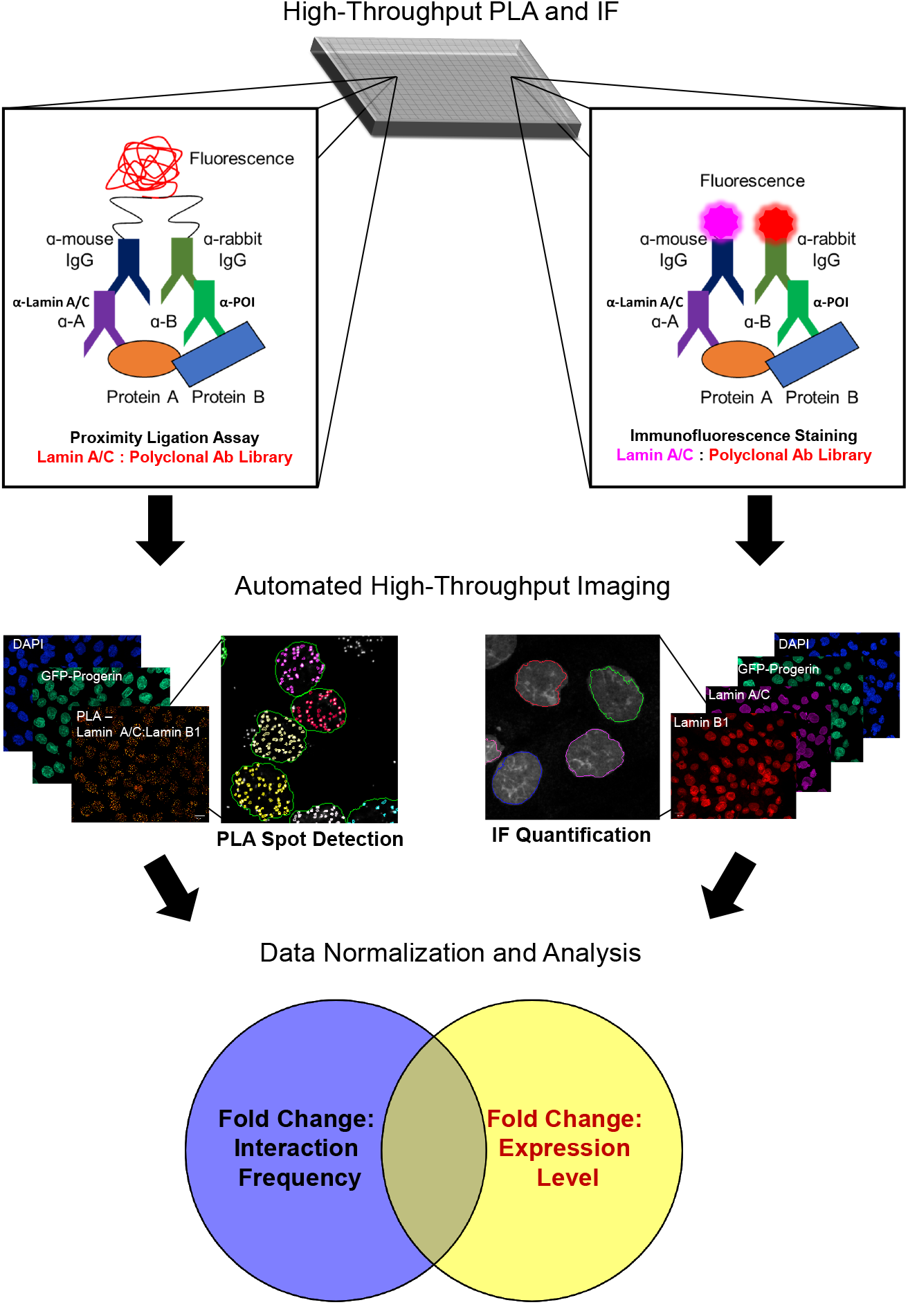
Outline of HiPLA. As a method to systematically interrogate changes to the protein interactome in response to a cellular intervention *in vivo*, the PLA was adapted for high-throughput imaging. PLA utilizes indirect antibody labeling to identify proteins of interest and secondary antibodies labeled with short DNA strands that are ligated when located within ~40 nm of each other. After ligation, rolling circle amplification with fluorescent oligonucleotides amplifies sites of antibody co-localization above the detection limit. As proof of concept, HiPLA is performed in 384-well plates on an immortalized fibroblast cell line with inducible expression of GFP-progerin to assay how the nuclear lamina interactome is affected in a model of HGPS. After a 48 h incubation in the presence or absence of doxycycline, cells are fixed, permeabilized, and blocked in 2% BSA. Using a library of antibodies against nuclear proteins, cells are either stained following the PLA protocol to identify interactions between a protein of interest with lamin A/C (left) or a conventional indirect IF protocol to label the protein of interest as well as lamin A/C (right). Following immunolabeling, cells are imaged in a high-throughput and automated manner. A representative example using lamin A/C and lamin B1 antibodies is shown. Under PLA-labeled conditions, image analysis is carried out to differentiate and to quantify fluorescent PLA foci in GFP-positive and -negative cells after creating a nuclear mask based on the DAPI image. For IF analysis, average nuclear mean fluorescence of the protein of interest as marked by the DAPI channel is quantified in GFP positive and negative cells. The relative changes in interaction frequency and interactor expression levels are then analyzed for changes in protein interaction frequency, interaction localization, and expression levels.

## 2. Methods

### 2.1. Cell culture

Generation and culture of immortalized fibroblasts with inducible expression of GFP-progerin has been previously described [26]. Briefly, hTERT immortalized dermal fibroblasts were generated after infection with lentivirus produced from both the doxycycline inducible pLentiCMVTRE3G-Neo-GFP-progerin plasmid (derived from Addgene #w813-1) and the constitutively tetracycline repressor A3 mutant expressing pLentiCMVrtTA3-Hygro plasmid (Addgene #26730). hTERT-TetOn-GFP-progerin cells were maintained in modified minimum essential medium (MEM, Thermofisher Scientific) containing 15% fetal bovine serum (FBS, Sigma Aldrich), 2 mM L-glutamine (Thermofisher Scientific), 100 U/mL penicillin (Thermofisher Scientific), and 100 μg/mL streptomycin (Thermofisher Scientific). To reach an optimal confluency of ~70%, approximately, 2,500 cells were seeded in each well of a clear bottom 384-well plate (PerkinElmer) in 50 μL of culture media in the presence or absence of 1 μg/mL doxycycline for 48 h.

### 2.2. Proximity ligation assay (PLA) and immunofluorescence (IF) staining

After 48 h of culturing, cells were washed twice in PBS and fixed in 4% paraformaldehyde (Electron Microscopy Sciences) for 10 min at room temperature. Cells were subsequently washed in PBS and permeabilized in 0.3% Triton X100 (Sigma Aldrich) for 10 min at room. Cells were again washed twice in PBS and either stored at 4 °C for staining at a later time or incubated with 2% BSA (Sigma Aldrich) for 1 h at room temperature. To conserve reagents, the subsequent reactions were performed in a total volume of 15 μL/well. Cells were incubated in primary antibody in 2% BSA. An optimized concentration of monoclonal lamin A/C antibody (1:250; clone E1, Santa Cruz Biotechnology) and the standard concentration to screen polyclonal antibodies (1:100) was determined using a known interaction as a positive control to ensure a resolvable, sub-saturating number of PLA foci would be obtained (see Results). All subsequent PLA reactions were performed using lamin A/C clone E1 antibody at a dilution of 1:250. After incubation with primary antibody for 1 h at room temperature, cells were washed 3 times for 10 min in either PBS for IF staining or 0.01 M Tris, 0.15 M NaCl, 0.05% Tween 20 (Buffer A) for PLA staining. For IF staining, cells were then incubated in Alexa Fluor 488-labeled donkey anti-mouse IgG secondary antibody (1:400; Thermofisher Scientific), Alexa Fluor 594-labeled donkey anti-rabbit IgG secondary antibody (1:400; Thermofisher Scientific), and 2 μg/mL DAPI for 1 h at room temperature. Cells were washed an additional 3 times with PBS prior to imaging.

In the case of PLA staining, cells were incubated in affinity purified donkey anti-mouse IgG Duolink In Situ PLA Probe MINUS (1:5; Sigma Aldrich) and affinity purified donkey anti-rabbit IgG Duolink In Situ PLA Probe PLUS (1:5; Sigma Aldrich) for 1 h at 37°C. After incubation, cells were washed 3 times for 10 min in Buffer A and incubated in 1 unit/μL T4 DNA ligase in diluted ligase buffer (1:5; Sigma Aldrich) for 30 min at 37 °C. Subsequently, cells were washed again 3 times 10 min in Buffer A and incubated in 10 units/uL DNA polymerase in diluted polymerase buffer with red fluorescence detection labeled oligonucleotides (1:5; Sigma Aldrich) and 2 μg/mL DAPI for 90 min at 37 °C. A final round of washes was performed twice for 10 min in 0.2 M Tris and 0.1 M NaCl then twice for 10 min in Buffer A.

### 2.3. High-throughput imaging and analysis

High-throughput confocal imaging was performed using a 63x water objective on a Yokagawa CV7000 spinning disk microscope at the CCR High-throughput Imaging Facility (NIH, Bethesda, MD, USA). For excitation, four laser lines were used (405, 488, 561, and 647 nm). Images were acquired with 0.5 μm Z-sections for at least 10 randomly chosen fields of view in an automated manner. Typically, at least 1000 cells were analyzed per condition. Images were processed using Columbus software (PerkinElmer). First, nuclei were identified and segmented using the DAPI nuclear stain. Nuclear PLA foci were identified using the built-in spot analysis script using DAPI-created masks and factoring in relative focus intensity as well as a splitting coefficient for accurate focus discrimination. An average of ≥2 PLA foci per nucleus was used as a cut-off for antibodies that showed no reaction based on a negative antibody control of lamin A/C clone E1 alone. For IF, the average mean fluorescence from the 561 nm channel present in the DAPI mask was used to determine relative levels and changes in protein expression. Average fluorescence obtained from the 488 nm excitation channel was used to filter progerin positive and negative cells.

### 2.4. Statistical analysis

After normalizing the average number of PLA foci per nucleus in progerin-expressing cells to wild type cells, a calculated Z-score of ≥ 2 or ≤ −2 was used to identify interactions sensitive to progerin expression. A calculated Z-score of ≥ 2 or ≤ −2 was similarly used to identify changes in average nuclear mean fluorescence. The replication protein RPA32, which did not interact with lamin A/C by PLA and is largely not associated with the nuclear lamina [27], was used as a control for changes in nuclear fluorescence intensity. Changes between antibody optimization conditions were analyzed using a one-way ANOVA.

## 3 Results

### 3.1. HiPLA methodology and calibration

Since its original inception to detail the levels of platelet-derived growth factor in solution [28], the PLA has been optimized for *in situ* detection of the interaction between two proteins of interest using antibodies of different originating species [8, 25]. This approach has been successfully utilized to probe a variety of cellular interactions including those of the lamin proteins with the polycomb complex [29], DNA damage foci [30], the E3 ubiquitin ligase Smurf2 [31, 32], and the protein phosphatase 2A protein CIP2A [33], as well lamin interactions with other nuclear membrane-associated proteins [34, 35]. Given the demonstrated utility of PLA in probing lamin function and the drastic changes observed in the organization of the nuclear lamina in the presence of the disease-causing progerin protein, we sought to develop the PLA in a high-throughput format, referred to as HiPLA, and applied it as proof-of-principle to comprehensively probe the lamin interactome in a model of HGPS.

HiPLA was routinely carried out in a 384-well format using a CV7000 spinning-disk confocal microscope combined with an automated image analysis pipeline for foci detection and IF quantification using PerkinElmer Columbus software (**Figure 1; see Methods**) [36]. At least 1000 cells were analyzed per sample. To initially optimize the assay, we tested 3 monoclonal antibodies against lamin A/C/progerin together with an antibody against the known lamin A-interacting protein lamin B1 as a positive control [37]. While all of the antibodies resulted in PLA foci, the lamin A/C clone E1 antibody showed the highest signal with an average of 79 ± 4.6 PLA foci per nucleus compared to 59 ± 3.6 and 58 ± 2.3 for the other two antibodies. Anti-lamin A/C clone E1 was therefore chosen for further optimization (**Figure 2A**). Negative control reactions performed in the absence of antibodies against lamin A/C or lamin B1 resulted in an average of less than 1 PLA focus per nucleus. Using serial dilutions of the lamin A/C and lamin B1 antibodies, an optimal antibody concentration was determined to maximize the number of PLA foci per nucleus. At a fixed dilution of 1:250 for anti-lamin B1, a dilution of 1:250 for anti-lamin A/C approached the maximum of 65 ± 8.3 PLA foci per nucleus (**Figure 2B**) (P < 0.0001 compared to the negative antibody control). Similarly, titration of the lamin B1 antibody against a fixed dilution of 1:250 for anti-lamin A/C showed a maximum of 60 ± 4.1 PLA foci per nucleus at a dilution of 1:100 (P < 0.0001 compared to the negative antibody control). A dose-dependent increase and saturation in PLA foci number upon antibody titration suggested that the assay was specific and the detected signals represented bona-fide protein interactions. Furthermore, we found no appreciable effect on PLA foci number as a function of reaction volume above 15 μL/well in 384-well plates (**Figure 2C**). Based on these optimization results, we performed our standard high-throughput assays using lamin A/C antibody clone E1 at a dilution of 1:250 and specific antibodies against potential interactors at a dilution of 1:100 in a reaction volume of 15 μL.

**Figure 2:**
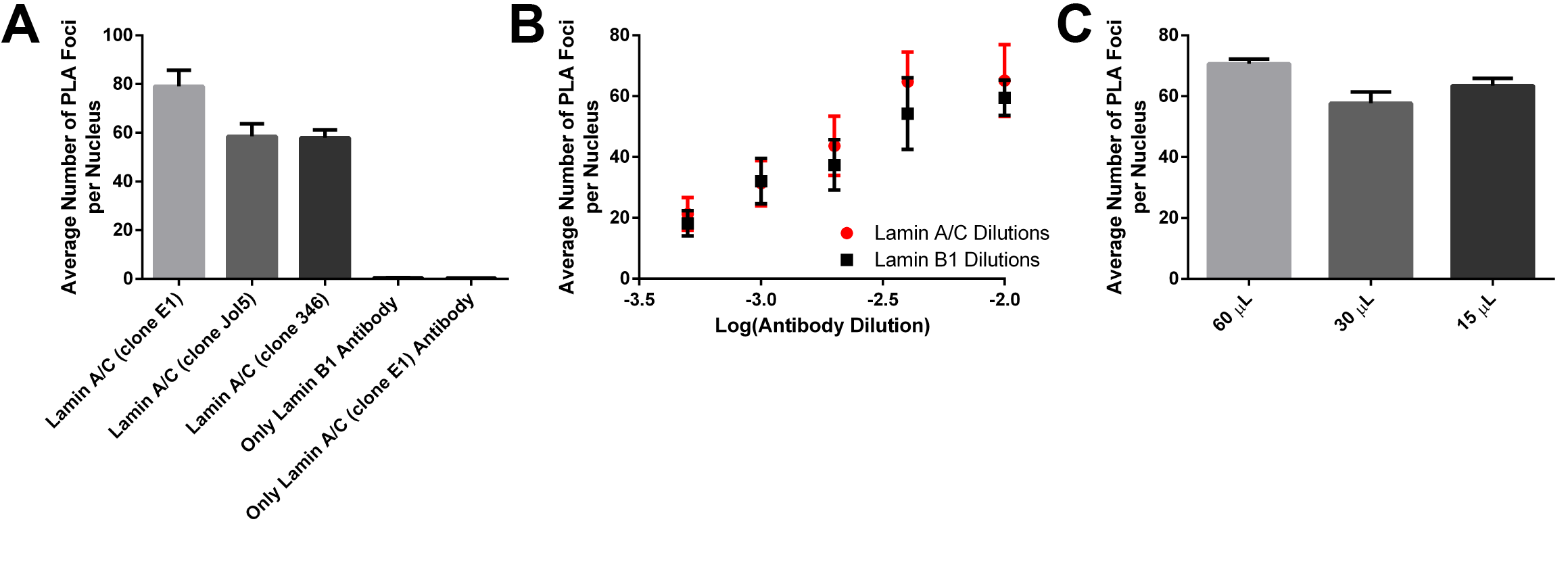
HiPLA optimization for lamin A/C interactors. (A) Antibody selection for HiPLA. HiPLA was carried out using a panel of antibodies to lamin A/C (1:100) in combination with an antibody to lamin B1 (1:100). While all three lamin A/C antibodies resulted in PLA foci as expected, the clone E1 antibody gave the largest average number of PLA foci per nucleus and was chosen for subsequent assays. (B) Antibody concentration optimization for HiPLA. PLA was performed using serial dilutions to determine the optimal antibody concentration of lamin A/C as well as an optimal working concentration for the panel of antibodies to potential lamina-interacting proteins. Dilutions were performed using a fixed concentration of anti-lamin B1 (1:250) and varying concentrations of anti-lamin A/C (red) or varying concentrations of anti-lamin B1 with a fixed concentration of anti-lamin A/C (1:250; black). (C) Reaction volume optimization for HiPLA. PLA using lamin A/C (1:250) and lamin B1 (1:100) antibodies were performed in decreasing reaction volumes to limit reagent consumption. No appreciable decrease in signal was observed below 15 μL/well. Values represent means ± SEM. N = 2 biological replicates and > 1000 cells per condition.

### 3.2. HiPLA of lamin A/C interactors in a HGPS model

We applied HiPLA to probe the lamin A interactome. Using the monoclonal lamin A/C clone E1 antibody, we screened the interactions of lamin A/C against a library of 133 polyclonal antibodies with a preference for nuclear proteins (**Supplementary Table 1**). 65 antibodies recognizing 45 distinct proteins and 15 specific post-translational modifications (PTMs) showed PLA signals above background levels (~49% of antibodies tested). In 7 cases where two antibodies against a protein or specific PTM were available, 5 proteins showed an interaction with both antibodies. However, because the absolute value of the PLA signal is highly dependent on the affinity and specificity of the antibody used, a lack of PLA signal does not exclude the possibility that an interaction does occur [23, 35].

Although the varying antibody quality precludes direct comparison of PLA signals between interactors, the relative PLA foci number using the same antibodies under different cellular conditions can be used to probe changes in interaction frequency of a given interaction pair. We therefore performed HiPLA in an immortalized skin fibroblast cell line expressing doxycycline inducible GFP-progerin [26] using the panel of 65 antibodies that had a positive PLA signal with lamin A/C (**Supplementary Table 1**). The change in interaction frequency for each antibody was assessed by plotting the ratio of the average number of PLA foci per nucleus in GFP-progerin expressing cells relative to wild type cells. Of the 60 lamin A/C interactors, 28 proteins and PTMs exhibited no change in PLA foci upon GFP-progerin expression and 32 proteins and PTMs showed a decrease by at least 2 standard deviations in the fold change of PLA foci per nucleus (**Figure 3**). Although only 1 of the two tested antibodies against H4K5ac showed a decrease in the fold change in PLA foci below 2 standard deviations, this interaction was still considered to be affected by progerin expression. Interestingly, none of the tested antibodies showed an increase in the fold change of PLA foci per nucleus by at least 2 standard deviations upon progerin expression. However, HES1, NCOR2, and XRCC4 did exhibit an increase in the number of PLA foci per nucleus albeit below the 2 standard deviation threshold routinely applied in our analysis. We conclude that HiPLA is able to successfully differentiate lamin A/C interactors and to effectively assay the effects on interaction frequency in response to progerin expression.

**Figure 3:**
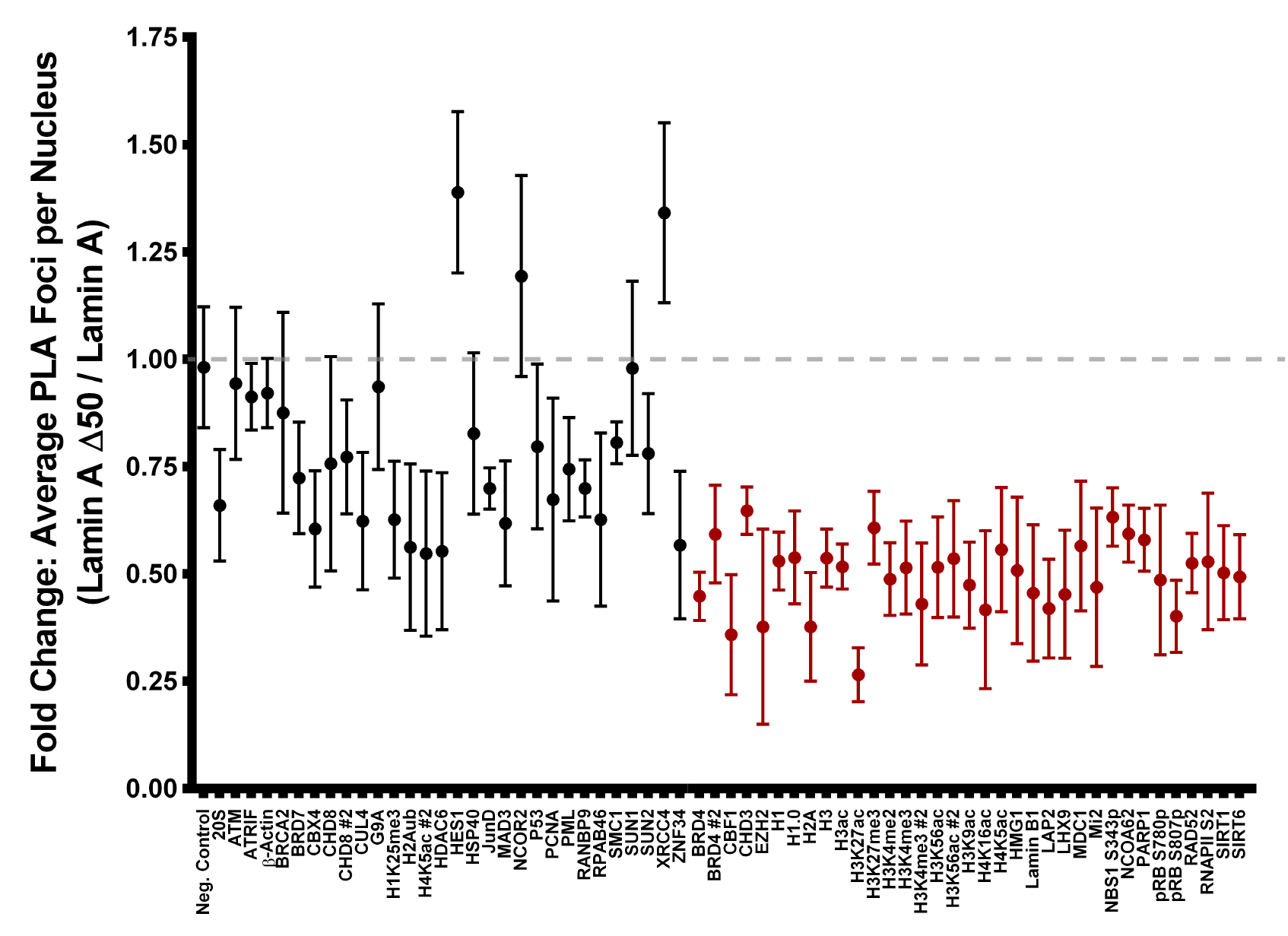
HiPLA screen for lamin A/C interactors in a model of HGPS. (A) HiPLA reveals changes in lamin A/C interactions upon progerin expression. HiPLA was performed on wildtype and GFP-progerin expressing cells with antibodies to lamin A/C clone E1 in combination with a panel of 65 antibodies of PLA-positive lamin A/C interactors. The average number of PLA foci per nucleus in GFP-progerin cells was plotted as fold change relative to wild type cells. Interactors that showed a >2-fold change in standard deviations of interaction frequency are shown in red. Gray dotted line marks an idealized ratio of 1, indicating no change in PLA foci number upon progerin expression. Values represent means ± SEM. N = 3–6 biological replicates and > 1000 cells per condition.

### 3.3. Combinatorial HiPLA and IF analysis in a HGPS model

While HiPLA was able to quantify the relative changes in the lamin A/C interactome, the assay does not provide information regarding the mechanism by which these interactions are affected. Detected interaction changes may either be due to direct (dis)association of the protein complex or due to changes in the level of the interacting protein which may indirectly affect complex formation. We therefore combined HiPLA with relative protein expression data obtained from quantitative high-throughput IF imaging (see Methods). High-throughput IF imaging was performed in the same manner as HiPLA with the exception that fluorophore-conjugated secondary antibodies were used for detection. Wild type and GFP-progerin expressing cells were imaged in 4 biological replicates and the fold change in nuclear mean IF intensity per antibody was determined. Of the 60 lamin A/C interactors, 29 proteins and PTMs were downregulated by at least 2 standard deviations and only SUN2 levels were upregulated by more than 2 standard deviations (**Figure 4A**).

**Figure 4:**
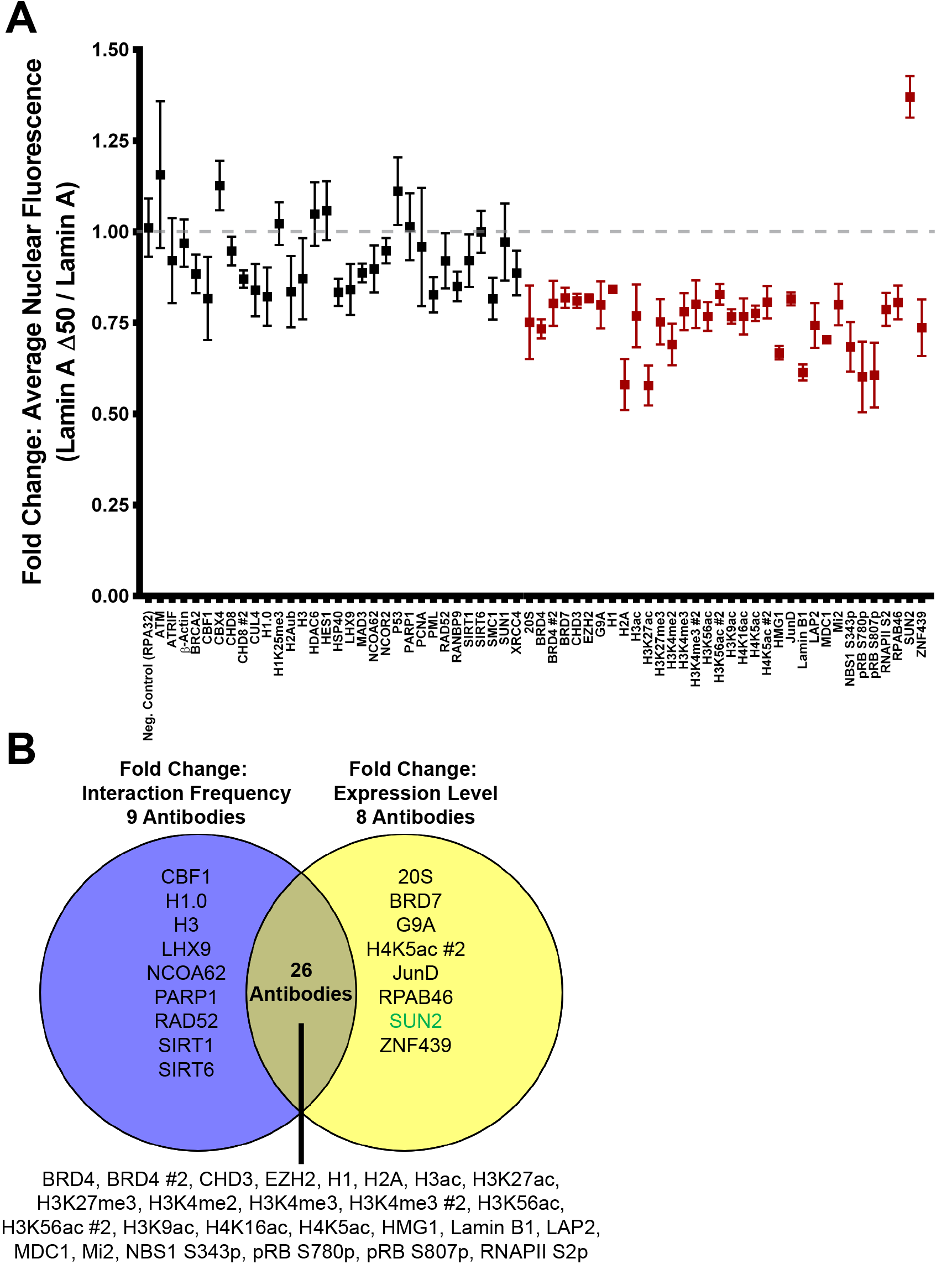
Combined HiPLA/IF to relate lamin A/C interactions to protein expression. (A) High-throughput IF reveals changes in lamin A/C interactor levels upon progerin expression. High-throughput IF was performed on wildtype and GFP-progerin expressing cells with antibodies to a panel of 65 antibodies of PLA-positive lamin A/C interactors. The average nuclear mean fluorescence intensity in GFP-progerin cells was plotted as fold change relative to wild type cells. Interactors that showed a >2-fold change in standard deviations of average nuclear mean fluorescence intensity are shown in red. Values represent means ± SEM. N = 4 biological replicates and > 1000 cells per condition. Gray dotted line marks an idealized ratio of 1, indicating no change in nuclear mean fluorescence intensity upon progerin expression. Fold change in RPA32 levels was used as a negative control. (B) Venn-diagram of lamin A/C interactors that showed a change >2-fold change in standard deviations of interaction frequency (average number of PLA foci per nucleus; from Figure 3; purple, left) and/or a >2-fold change in standard deviations of expression (average nuclear mean fluorescence intensity; from (A); yellow, right). Not shown are the 21 lamin A/C interactors that did not exhibit a change in interaction frequency nor expression. Green text indicates an increase; black text indicates a decrease.

We finally asked whether the decreased interactions identified by HiPLA were correlated with changes in protein expression or whether the changes in protein complex formation occurred independent of interactor expression level. Of the 65 antibodies that interacted with anti-lamin A/C, we found 22 antibodies against 21 proteins that showed no change in either interaction frequency or expression (**Figures 3, 4A**). 26 antibodies against 23 proteins and PTMs exhibited both a decrease in the fold change in the average number of PLA foci per nucleus and in the average mean fluorescence intensity upon progerin expression (**Figure 4B**). Thus, corresponding changes in interaction frequency and expression were observed in 48 of 65 antibodies (73%). A change in protein expression without a corresponding change in interaction frequency was observed for 20S, BRD7, G9A, JunD, RPAB46, SUN2, ZNF439, as well as for one of the H4K5ac antibodies (**Figure 4B**). Alternatively, a decrease in interaction frequency without a corresponding decrease in expression was observed for CBF1, H1.0, H3, LHX9, NCOA62/SKIP, PARP1, Rad52, SIRT1, and SIRT6 (**Figure 4B**), suggesting that these interactors may be affected by progerin via direct inhibition of complex formation. Together, HiPLA with high-throughput IF imaging systematically identified changes in protein interactions and correlated these changes to potential mechanisms of disease.

## 4. Discussion

We have developed HiPLA, a high-throughput version of the well-established proximity ligation assay for the detection of protein-protein interactions *in vivo*. Using HiPLA, we have identified a number of lamin A/C interactors and investigated how these interactions are modified in a cellular model of the premature aging disorder HGPS. In combination with high-throughput IF staining, we demonstrated that the majority of the observed changes in lamin interactions are paralleled by changes in the expression of the interaction partners. This exploratory screen showcases the benefit of multiplexing IF and PLA immunoassays in a high-throughput pipeline to discern the localization and frequency of protein complex formation.

We have identified 65 antibodies against 60 proteins and PTMs that react with lamin A/C by HiPLA. Many of these proteins have previously been described as lamin A/C interactors by mass spectrometry analysis [5, 20, 21, 38], demonstrating the specificity of our assay (**Supplementary Table 1**). Intriguingly, we identified several lamin A/C interactors including transcription factors JunD, HES1, LHX9, and MAD3 as well as the DNA damage response components ATM, NBS1, RAD52, and XRCC4, which to our knowledge have not been previously explored as lamin A/C interactors. Of the 65 antibodies that resulted in a PLA signal with lamin A/C, 35 antibodies against 32 proteins and PTMs exhibited a decrease in interaction frequency upon progerin expression. This finding is in line with the large scale changes in gene expression identified in models of HGPS [39- 44] well as differences in the lamin A/C interactome previously reported by mass spectrometry analyses [21, 22, 38].

In combination with high-throughput IF, we noted a strong correlation between PLA signals and expression level of lamin A/C interactors, suggesting that the reduced interactions in progerin-expressing cells may be a consequence of limited protein availability. However, CBF1, H1.0, H3, LHX9, NCOA62/SKIP, PARP1, Rad52, SIRT1, and SIRT6 exhibited reduced interactions with the lamina without reductions in protein level. While the association of lamin A/C with LHX9 or Rad52 has not previously been investigated, lamin A has been shown to both directly and indirectly interact with histones, and this interaction may be impaired in HGPS [12, 45-48]. Additionally, a previous study reported that lamin A, but not progerin, is able to bind to and activate SIRT1 [49]. Lamin A is also necessary for SIRT6-mediated activation of PARP1 ribosylation upon DNA damage [50]. Unlike SIRT1, both recombinant or transiently over-expressed lamin A and progerin were found to bind SIRT6 by co-immunopercipitation, however, only lamin A was able to stimulate SIRT6 deacetylase activity [50]. Although further studies are needed to reconcile this finding with our observation that the association between SIRT6, PAPR1, and the lamina is decreased upon progerin expression, the impaired ability of inactive SIRT6 to bind chromatin [50, 51] may also destabilize its interaction with the nuclear periphery. Furthermore, as observed here, progerin expression has been reported to contribute to the activation of Notch target genes by loss of localization of the transcriptional co-activator NCOA62/SKIP at the nuclear periphery [40]. While we did not observe a change in NCOA62/SKIP or NCOR2 expression as previously reported [40], we did find that the interaction between CBF1, which associates with NCOA62/SKIP [52], and the lamina was impaired, suggesting progerin-expression may also cause other Notch signaling factors to differentially interact with the nuclear periphery.

Our results demonstrate that HiPLA is a versatile method to identify the effects of cellular interventions on protein complex formation. While PLA has been found to be less sensitive than FRET based protein-protein interaction detection [53], our ability to confirm reported changes in protein complex formation with progerin expression such as H3K27me3 [45], lamin B1 [54], LAP2 [22], and SIRT1 [49] suggest that PLA is well-suited for detection of robust changes in interaction frequency. In addition, PLA is well suited for scaling up to a high-throughput format to analyze tens and hundreds of potential interactions, which has been challenging for other fluorescence-based methods. Although PLA also offers information about the localization of interactions, we found that fluorescent PLA foci were too large to accurately discriminate interactions at the nuclear periphery from the nucleoplasm and this was further complicated by the nuclear invaginations induced by progerin expression (data not shown). Optimization of the method using shorter amplification times, cells with larger nuclei, co-staining with markers of nuclear compartments, and thinner imaging sections may afford better resolution in the Z-plane to accurately localize interactions within the nucleus. Nevertheless, HiPLA does seem ideally suited to assay interactions between proteins that form complexes in different parts of the cell.

HiPLA also has the potential to be combined with other fluorescent assays to expand its utility. Because the PLA detection of an interaction between two proteins only uses one imaging channel, IF for the interacting proteins can be imaged simultaneously in the same well if a monoclonal antibody with a non-overlapping epitope or polyclonal antibodies are employed. IF imaging can be used to correlate interaction data with expression levels, as done here, as well as to relate the site of interaction to cellular landmarks using localization markers or to the distribution of other protein complex markers. Furthermore, if complemented with siRNA knockdown, CRISPR knockout, or drug screening libraries, this methodology has the potential for large-scale screening for mediators of protein-protein interactions. Taken together, HiPLA is a high-content imaging method that can be customized to comprehensively probe how cellular interventions influence the protein interactome and will be useful in the discovery of regulators of protein-protein interactions.

## 5. Acknowledgments

This research was supported by the Intramural Research Program of the National Institutes of Health, National Cancer Institute, and Center for Cancer Research. High-throughput imaging work was performed at the High-Throughput Imaging Facility (HiTIF)/Center for Cancer Research/National Cancer Institute/NIH.

---

**Supplementary Table 1.**
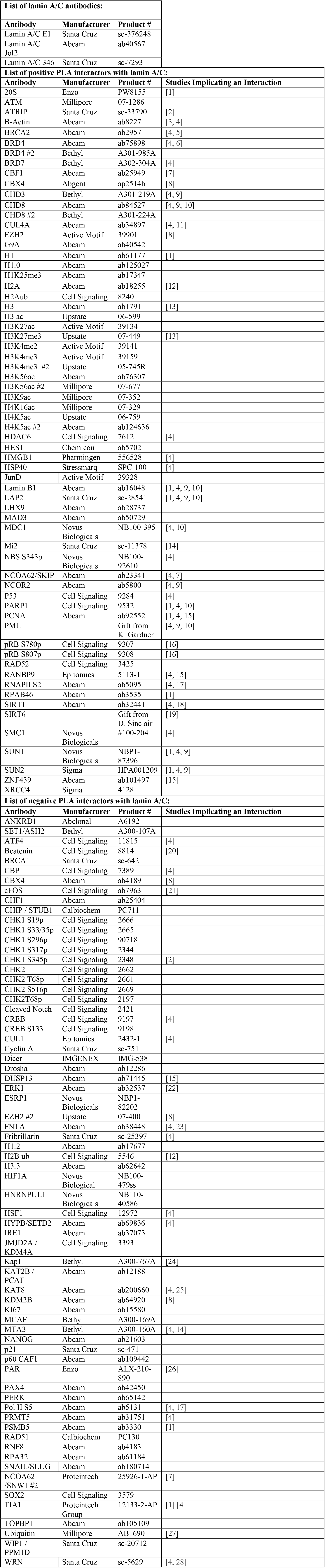

